# Concept and in-silico assessment of an algorithm for monitoring cytosolic fluorescent aggregates in cells

**DOI:** 10.1101/177139

**Authors:** Yasel Garcés Suárez, Vadim Pérez Koldenkova, Tomoki Matsuda, Adán Guerrero, Takeharu Nagai

## Abstract

Autophagy is an evolutionary conserved pathway, by which eukaryotic cells degrade long-living cellular proteins and intracellular organelles, to maintain a pool of available nutrients. Impaired autophagy has been associated to important pathophysiological conditions, and this is the reason why several techniques have been developed for its correct assessment and monitoring. Fluorescence microscopy is one of these tools, which relies on the detection of specific fluorescence changes of targeted GFP-based reporters in dot-like organelles in which autophagy is executed. Currently, several procedures exist to count and segment this punctate structures in the resulting fluorescence images, however, they are either based on subjective criteria, or no information is available related to them. Here we present the concept of an algorithm for a semi-automatic detection and segmentation in 2D fluorescence images of spot-like structures similar to those observed under induction of autophagy. By evaluating the algorithm on more than 20000 simulated images of cells containing a variable number of punctate structures of different sizes and different levels of applied noise, we demonstrate its high robustness of puncta detection, even on a high noise background. We further demonstrate this feature of our algorithm by testing it in experimental conditions of a high non-specific background signal. We conclude that our algorithm is a suitable tool to be tested in biologically-relevant contexts.

## Introduction

Autophagy is an evolutionary conserved pathway of degradation of long-lived cellular proteins and organelles, to recycle valuable cellular nutrients. Mechanistically, it involves the enwrapping of cellular cytosolic material in the so-called autophagosomes, which is then delivered to lysosomes, so the content of the resulting organelles — autolysosomes — is degraded to be reused [1]. There is always a basal level of autophagy in nutrient-replete conditions, but under stimuli like starvation, autophagy is upregulated, to maintain the intracellular pool of aminoacids and other vital compounds [2].

Albeit its apparent role as a simple degradation system, autophagy is required to prevent neurodegeneration, for suppression of tumors, for the regulation of the immune system, for elimination of intracellular microbes and alleviation of aging effects ([1] and references therein). Given the importance autophagy has in the prevention of diseases, much effort have been dedicated to the establishment of methods for its correct identification [1, 3]. As autophagy is a dynamic process, its main characteristic is the rate of the steps constituting it, being all them collectively known under the term “autophagic flux” [1, 4]. Among the existing techniques for monitoring this flux, fluorescence microscopy is probably the only one which allows following it in real time. For this purpose, green fluorescent protein, GFP, is employed, specifically targeted to autophagosomes by its fusion with the microtubule-associated protein 1 light chain 3 (LC3) [5]. Formation of autophagosomes can be visualized in this case as an increase of GFP-positive punctate structures in the cell cytosol. However, a drawback of this system is that autophagy endpoint cannot be distinguished, as GFP fluorescence is quenched in the acidic environment following fusion of autophagomes with lysosomes. To overcome this problem an improved autophagy indicator has been proposed, which in addition to GFP includes a less-acid sensitive red-emitting fluorescent protein, which signal can be observed in autolysosomes even after the disappearance of that one from GFP [6].

The analysis of the images obtained with fluorescent protein-based indicators requires a reliable procedure for the segmentation and counting of cytosolic fluorescent spots. This step is typically performed either manually, using commercially available tools (Top Hat of MetaMorph, or G-Count by G-Angstrom, [1]), or the ImageJ plugin WatershedCounting3D [7]. Unfortunately, information on how is segmentation carried out could not be found (G-Count, WatershedCounting3D), or is subjective and depends on the person conducting this analysis (manual analysis and the Top Hat algorithm). Moreover, we could not find evaluations of the segmentation accuracy of every of these algorithms.

Here we propose an algorithm for the identification of autophagosome/autolysosome-like cytosolic spots using a dynamically-calculated threshold value. Using more than 20 000 simulated images with different noise levels and variable number and size of spots, we analyzed the performance of the algorithm, and show the robustness of meaningful signal detection even in conditions of very high background noise. To further test the algorithm performance, we applied it to analyze images obtained in experiments with cells which expressed a tandem indicator in the cytosol, i.e. displaying a higher non-specific cytosolic signal, that in conditions when the indicator is directly targeted to the autophagosomes. Two-three days after transfection cells expressing the cytosolic indicator displayed fluorescent punctate structures which were successfully detected and segmented by the algorithm, even on a background of high cytosolic fluorescence.

## 1 Materials and methods

### 1.1 Plasmids construction

Construction was carried in the pRSETB vector (Invitrogen, USA). Amplified and digested mCherry and cp173Venus sequences were cloned in the named vector between *Bam*HI and *Eco*RI sites through an intermediate *Xho*I site using three-piece ligation. For expression in mammalian cells, the mCherry-cp173Venus sequence was subcloned into the pcDNA3.2 vector (Life Technologies, USA). Plasmids propagation was performed in the XL10 Gold *Escherichia coli* line, and purification for mammalian cell transfection was performed using a PEG-8000-based largeprep protocol.

### 1.2 Cell culture and transfection

HeLa cells (from RIKEN BioResource Center, Japan) were cultured at 37*°* C in a 5% CO_2_ atmosphere in DMEM medium (Sigma Aldrich) +10% heat-inactivated FBS (Biowest, Canada). Cells growing in 100-mm plastic dishes were passed twice a week with medium replacement. For imaging experiments the cells were additionally plated on Cellmatrix (Nitta Zeratinn, Japan)-coated custom-made glass bottom dishes, filled with 2 mL of medium of the same composition.

HeLa transfection was performed directly in the glass bottom dishes 24 hrs after plating, following a variant of the “Ca^2+^-phosphate protocol” [11]. Transfected cells were then cultured for another 48-72 hrs until imaged.

### 1.3 Imaging and cell stimulation

Imaging was performed at room temperature with a Nikon Ti Eclipse confocal microscope (through a 60 × Plan Apo VC oil-immersion objective lens [NA: 1.40]), equipped with an A1 imaging system. To avoid bleeding of the Venus emission into the mCherry channel their excitation was performed sequentially using the built-in laser lines (488 nm for cp173Venus [excitation max — 516 nm], and 561 nm for mCherry [excitation max — 587 nm]). Emissions were read using the manufacturer-provided emission filters in the following ranges: 500-550 nm (cp173Venus; emission max — 530 nm) and 570-620 nm (mCherry; emission max — 610 nm). Individual images were acquired with 5-sec intervals; the total duration of imaging was as specified in the corresponding figures, constituting 3 to 10 mins. Perfect Focus system was enabled during imaging to avoid defocusing of the sample. For imaging, glass-bottom dishes were directly transferred from the incubator to the microscope stage.

### 1.4 Parameter selection of simulated images

Simulated images were set to contain from 20 to 50 fluorescent spots of sizes between 0.1 and 1.2 *μm*, similar to data shown in [1, 4, 8].

The simulated images were generated using the approach proposed by Sinko and collaborators in [9]. The next algorithm resume the steps for the simulated images generation:

#### Algorithm 1: Simulated Image Generation

1. Creation of image *A* containing fluorescent organelles (simulated as little ellipses) of biologically-relevant sizes and positions. The number of the organelles, the size and positions are generated randomly using a uniform distribution function.

2. Convolute *A* using the point spread function (PSF) [10]. For this step we used the plug-in “Diffraction PSF-3D” of ImageJ.

3. Add Gaussian noise in order to consider auto-fluorescence and Poisson noise for the purpose of considering electronic noise.

4. Finally, the image is multiplied by 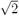 to consider amplification in a EM-CCD detector [9].

In the step 2 of the Algorithm 1 we consider the following parameters for the image convolution: Index of refraction of the media *n* = 1.33, Numerical Aperture *N A* = *n* sin(*θ*) = 1.3, Wavelength *λ* = 510 *nm*, Image Pixel Spacing *px* = 100 *nm*, Width (pixels) = 512, Height (pixels) = 512. Rayleigh resolution *r* = 0.61*λ/N A* = 240 *nm*. For more details of this plug-in see [10].

In Fig. 1 the steps of the Algorithm 1 for the generation of simulated images is visually represented. The Fig. 1(d) can be considered as an example of the images that were used to the validation of the proposed algorithm.

**Fig 1.**
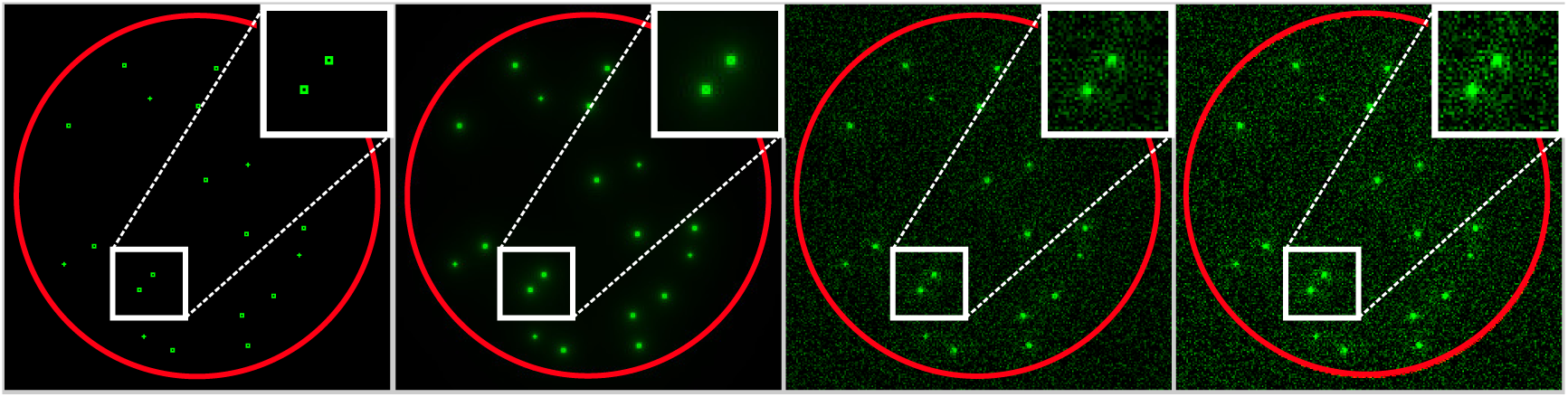
Visual example of the simulated images generation algorithm. **(a)** Generation of the organelles. **(b)** Convolution of the image (a) using the point spread function. **(c)** Image contamination with Gaussian and Poisson noise. **(d)** Image with the simulated amplification in a EM-CCD detector. The red circumference simulate the cell and only was included for a more accurate visual reference.

### 1.5 Image analysis

Detection and quantification of cytosolic puncta in cells was performed semi-automatically in two steps:

#### Manual mask creation

For an each image sequence *ℐ*, a mask image *M* _*ℐ*_ *∈ M* (*m, n*) is created manually. This step allows to isolate each cell {*C*_1_*, C*_2_*, …, C*_*K*_} from others, then *M*_*I*_ is defined as:

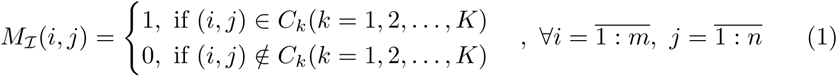

where *M* (*m, n*) is the image set of dimension *m × n*.

#### Puncta detection

For a cell *C*_*k*_ *∈* {*C*_1_*, C*_2_*, …, C*_*K*_} of the image sequence *I* in a time *t*, we denote this cell as *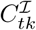* and then perform the next steps:

1. *Filter the cell* : This step consists in thresholding the image *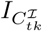* (denotes the image that contain only the cell *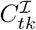*) to remove noise and atypical (low-intensity) values. The threshold (𝔍) is computed dynamically in dependence of the characteristics of each images, and as consequence, offers an adaptive way of elimination of noise and atypical values for each images (a detailed explanation is available in Section 1.5.1). The filtered image is defined as:

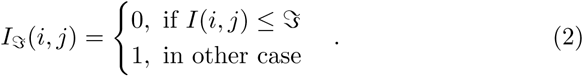
2. *Compute the connected components*: The connected components labeling is used to detect connected pixels, corresponding to a single lysosome/puncta, in the image *I*_*𝔍*_ (*i, j*). For this, each pixel of the image is visited, and then external boundaries are created using pixel neighbors according to a specific type of connectivity [12]. The results of this step are *N* connected components *A*_*i*_*, i* = 1, 2*, …, N*, where each *A*_*i*_ is identified as a single lysosome/puncta. In this step it is assumed that cytosolic fluorescent aggregates are composed of several pixels, and a digital path exists between them.

##### 1.5.1 Threshold by entropy

From the point of view of digital image processing, the entropy corresponds to the average of the information contained in the image, and is defined as:

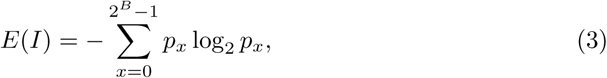

where *B* is the total number of bits of the digitized image *I* and 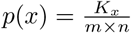 is the probability of occurrence of a gray-level value, *K*_*x*_ is the number of times that the pixel with a value *x* is observed in *I*, *m* and *n* are the numbers of rows and columns of the image respectively, and as consequence *m* × *n* is the total number of pixels in the image. The minimum of entropy is achieved when the image itself is constant (all pixels have the same value) and in this case *E*(*I*) = log_2_ 1 = 0. The maximum entropy is reached when the pixel’s values follow a uniform probability distribution function, this is, if *B* is the number of bits by pixel, then exists 2^*B*^ gray values in the image and the probability of each of them is exactly:

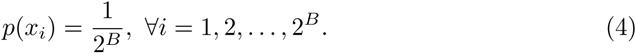

If we consider the definition of entropy (3) and the condition (4), then:

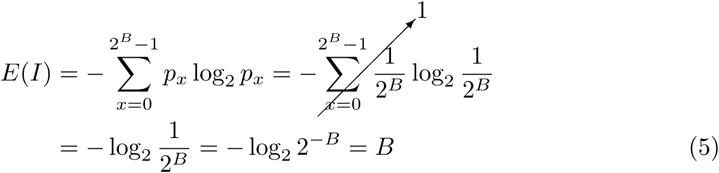

The above information provides the limits for the entropy function of a digital image of *B* bits per pixel:

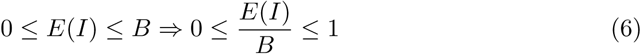

Considering the entropy function as a measure of the information contained in an image, it is possible to interpret this value as an index that allows to compute the relationship between the signal and the noise in the image. For example, low entropy images, such as those containing a limited number of small biological structures, have very little contrast and a large number of pixels with the same or similar grays values (not much signal). On the other hand, high entropy images such as those with a large number of fluorescent biological objects, have a large contrast between adjacent pixels.

#### Definition 1 (Entropy Information Criterion (EIC)) *The entropy information criterion is defined as:*

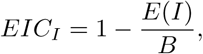

*where E*(*I*) *denote the entropy of the image I and B is the number of bits per pixels in I.*

Note that *E*(*I*)*/B ∈* [0, 1] (see equation (6)) and as consequence *EIC*_*I*_ *∈* [0, 1]. The scalar *EIC*_*I*_ could be interpreted as a measure of the percent of redundant (noise) information in the image.

#### Percentile Computation

For a serie of measurements {*Y*_1_*, Y*_2_*, …, Y*_*N*_} denote the data ordered in increasing order. The p-th percentile is a value *Y* (*p*), such as (100 × *p*)% of the measurements are lower than it, and at most (100 *×* (1 -*p*))% are greater [13].

The p-th percentil is obtained by calculating the ordinal rank, and then taking from the ordered list the value that corresponds to that rank. The ordinal rank *n* is calculated using the formula:

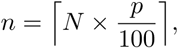

where *N* are the total number of observations, *p ∈* [0, 1] is the percent of the data less than *Y* (*p*). The operator ⌈ *·* ⌈ is the integer part, and is defined as:

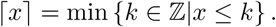

The value from the ordered list that corresponds to the ordinal rank is the percentile, that is, *Y* (*p*) = *Y*_*n*_.

The combination of the EIC (see Definition 1) and the computation of the percentile provide a clear and easy way to determine the threshold of any image.

#### Definition 2 (Entropy Threshold Criterion (ETC))

*Let I be an image with a colour depth of B bits per pixel, the Entropy Threshold Criterion of I is defined as:*

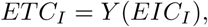

*where EIC*_*I*_ *is the “Entropy Information Criterion” defined in Definition 1 and Y* (*·*) *is the EIC*_*I*_ *percentile of the image pixels values.*

The ETC provides a fast, simple and efficient alternative to compute the threshold value for an image of any size and resolution. Also, it can be easily implemented in any programming language and allows to calculate in a dynamic way the threshold value for image sequences.

Applying this procedure to fluorescence imaging, we can automatically select in the image those pixels with the highest intensity values. The threshold value 𝔍 defined in the equation (2) is computed using the Definition 2.

### 1.6 Evaluation of Signal/Noise ratios (SNR)

In the analysis of the performance of algorithm, it is important to take into account the relationship between the level of signal versus noise power. According to Gonzalez and Woods [14], if we denote by *R*(*x, y*) the image that only contains the signal and by *C*(*x, y*) the corrupted image (containing both signal and noise), then, the signal/noise ratios (SNR) can be computed as:

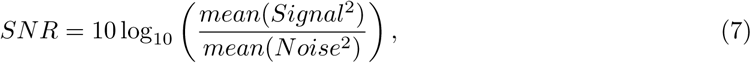

Where

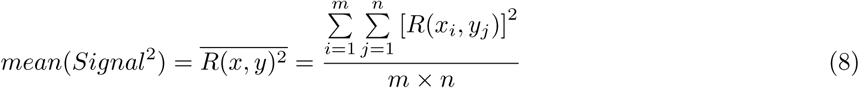

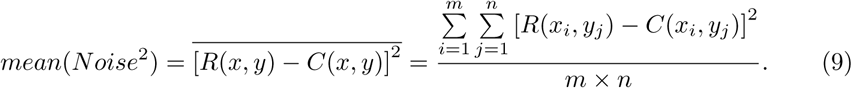

Then, replacing (8) and (9) in the equation (7) we obtain:

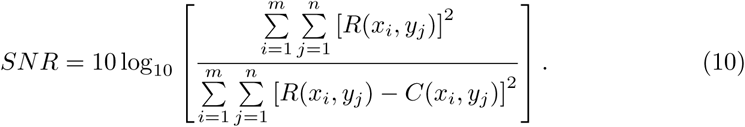

The images *R*(*x, y*) and *C*(*x, y*) have a size of *m* × *n* pixels. The SNR is expressed in decibels (dB) and higher numbers correspond to better contrast, since there is more useful information (the signal) than unwanted data (the noise). For example, when an image have a *SN R* = 2*dB*, it means that the signal is 2 times higher than the level of the noise. Note that with this definition of SNR it is possible to obtain negative values of SNR, this situation occurs when the image contains more noise than signal.

## 3 Results and Discussion

### 2.1 Accuracy of the segmentation algorithm

The images obtained as a result of the process contain fluorescing organelles of known sizes and positions. This facilitates the analysis of the algorithm’s response against several plausible events, hence permitting comparison of the obtained results with “ground truth” organelles considering different signal/noise ratios.

Let 𝕃_*I*_ = *{ℒ*_*i*_*, i* = 1, 2*, …, n}* and 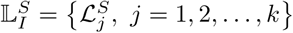 be the set of organelles that has been generated in the simulated image I, and the set of organelles obtained as a result of the segmentation algorithm respectively. The organelles in both sets are uniquely identified by their centers (*x, y*), and as consequence the sets 𝕃_*I*_ and 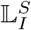 can be compared following the next criteria:

#### True Positive (TP)

The organelle spot *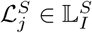* is a true possitive if:

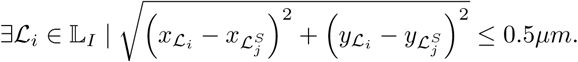

#### False Positive (FP)

The organelle spot *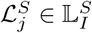* is a false possitive if:

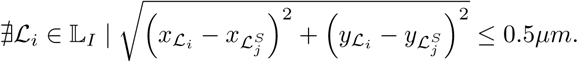

#### False Negative (FN)

The organelle spot *ℒ*_*j*_ ∊ *𝕃*_*I*_ is a false negative if:

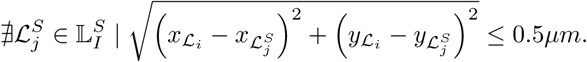

In each of the previous definitions 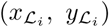 and 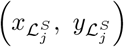 denote the center of mass of the organelles *ℒ*_*i*_ *∈* 𝕃_*I*_ and 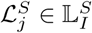 respectively. Note that in each case 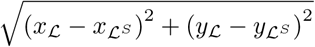 is the euclidean distance between the center of mass of the organelles *ℒ*_*i*_ and *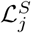*

Similarly to the work of Garcés and collaborators [15], the 0.5*μm* threshold was chosen based on the resolution limit of optical microscopes, which in agreement with Rayleigh’s criteria, is *r* = (0.61*λ*)*/N A*, with *λ* being the emission wavelength of the fluorophore and *N A* the numerical aperture of the objective. For the sake of simplicity, we define a 0.5*μm* threshold as an approximation to the diameter of the zero-order Airy ring, because in diffraction-limited images it does not make sense to segment structures smaller than 2*r*.

The Fig. 2 represents the criteria described previously. A TP is returned when an organelle is detected within a distance shorter than 0.5*μm* to that of a simulated spot. A FP result is returned when the algorithm detects an organelle that does not exist in the “ground truth” image. A FN is obtained when a organelles in the “ground truth” images is not detected by the algorithm.

**Fig 2.**
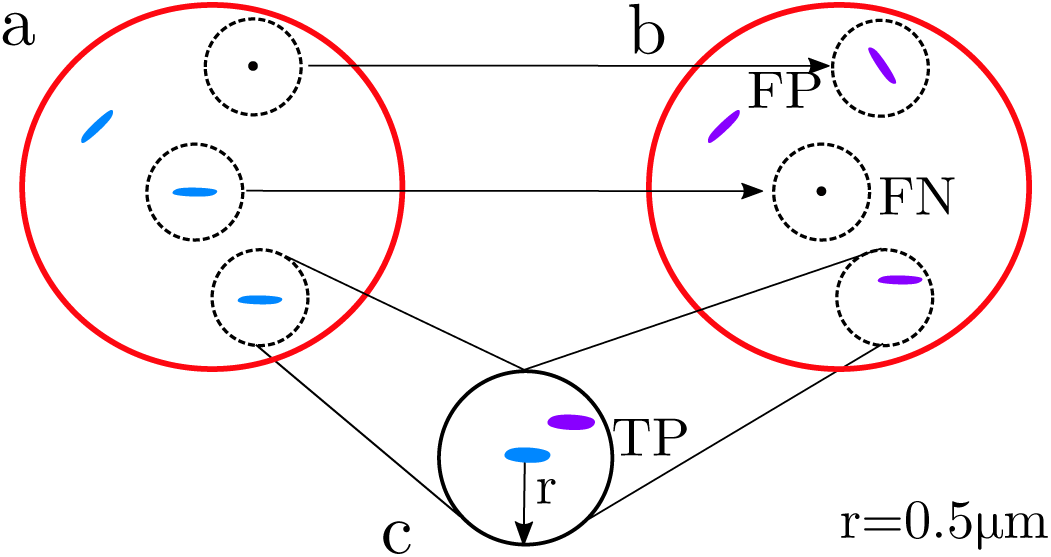
Criteria of True Positive (TP), False Positive (FP) and False Negative (FN) criteria. **(a)** Representation of the “ground truth” image. **(b)** Result of the segmentation algorithm. **(c)** Zoom of a TP classification. The red circumference simulate the cell and all the dashed black circumferences inside the cell represent the ROI and have a radio of 0.5*μm*.

The criteria presented above allowed to carry up the validation of the algorithm using indexes like the Jaccard index, the Recall and the Precision.

The Jaccard index [16] is a measure of similarity for two sets of data, with a range from 0 to 1. The higher the value, the more similar the two sets are, and as consequence, is very easy to interpret. If we consider the sets 𝕃 _*I*_ and 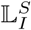, the Jaccard index is defined as:

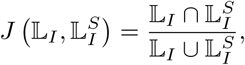

in our specific case, and considering the criteria presented before, we obtain that:

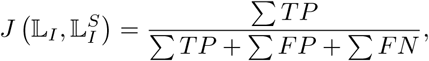

where Σ *T P, Σ F P* and *F N* indicate the total number of true positive, false positive and false negative respectively.

The Precision [17] (positive predictive value) is the fraction of organelles that were well detected (TP) among all the organelles found by the algorithm (TP+FP). It is defined as:

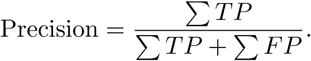

The Recall (true positive rate or sensitivity) [17] in the context of our application is the fraction of organelles that have been detected by the algorithm over the real number of organelles in the “ground truth” image. It is defined as:

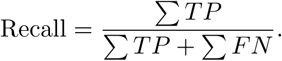

The Precision and Recall take values in the interval [0, 1]. The higher values indicate a better accuracy of the algorithm.

The performance of the algorithm was evaluated on more than 20 000 simulated “ground truth” images generated taking into account different SNR values, number, size and position of the organelles. In the simulated images we included high levels of noise (see step 2 in the Algorithm 1) with the intention to observe the stability and performance of the algorithm in extremely complex scenarios.

Fig. 3 shows the obtained results for the Precision, Recall and Jaccard indices as a function of the SNR. The curve of the Jaccard’s index show that even for extremelly noisy images (*SN R* ≤ 0*dB*) the algorithm have a good performance with values, in all the cases, over 0.7 (over 70% match between the sets 𝕃_*I*_ and 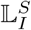). A very similar behavior is observed in the case of Precision, where the index value is directly proportional to the SNR, note that even when the *SN R* = -8*dB*, more than 70% of the organelles detected by the algorithm correspond to organelles in the “ground truth” images. The Recall have values over 0.97 for all the SNR, this means that, approximately 97% of the organelles in the “ground truth” images were detected by the algorithm for all the SNR. In general, the algorithm show a high robustness to noise, even in cases when noise is much higher than the signal in the images (*SN R* ≤ 0*dB*). The values obtained show that our procedure offers a very good detection level of the organelles in the original images.

**Fig 3.**
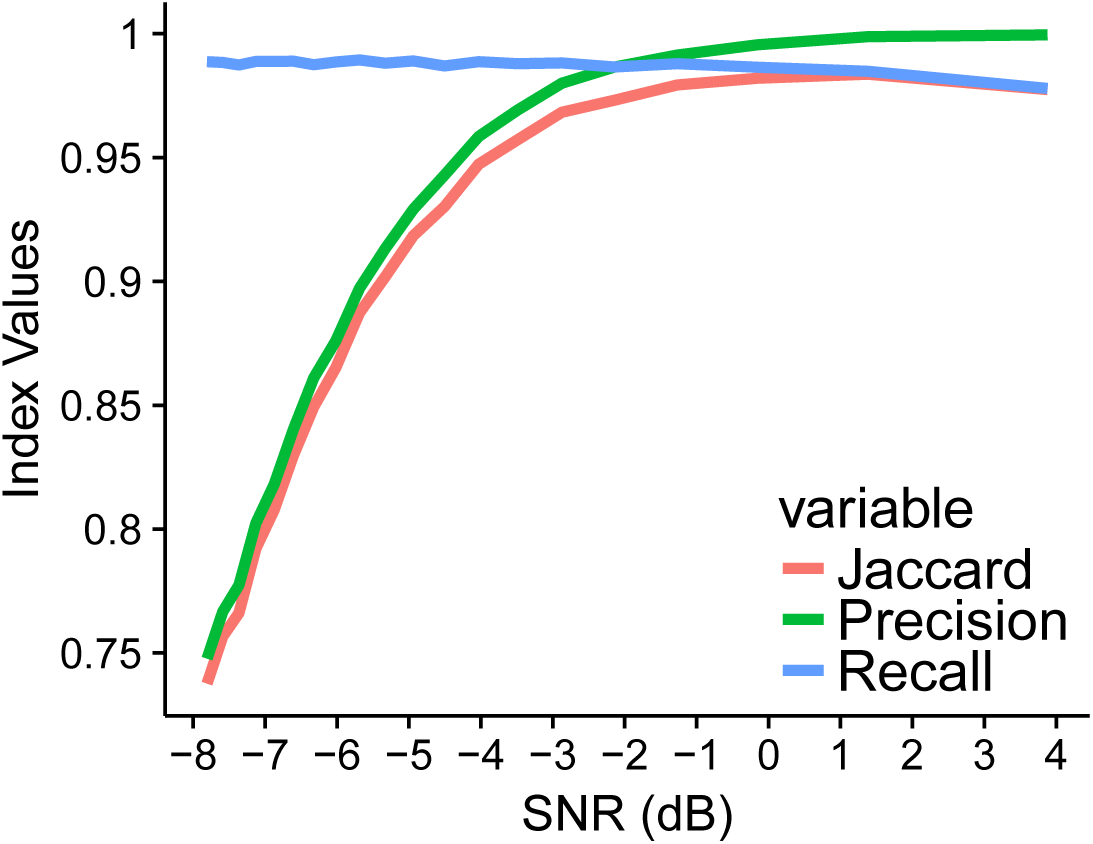
Plots of Precision, Recall and Jaccard indexes for the detection and adjustment of the organelles considering different levels of SNR in the simulated images.

One of the advantages of the algorithm is the possibility to compute some properties like the area, the mean intensity, the perimeter and the spatial position of each detected spot corresponding to an organelle. We use the information about the area of each “ground truth” organelle to study the error in the adjustment of each organelle, so, we can use this information to analyze the performance of the algorithm in fitting the real size of a organelle.

**Note 1** *If 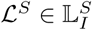 is a true positive detected organelle, then by definition (see the criteria above):*

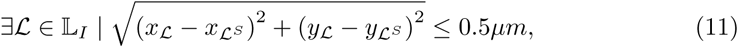

*then it is possible to compare the area of the organelles ℒ*_*i*_ *and ℒ^S^. The error in the area of the organelles ℒ and ℒ^S^ is defined as:*

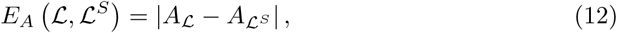

*where |·| denote the absolute value.*

Fig. 4(a) shows the boxplots for the error in the area fitting of the organelles (see equation (12)) in function of the SNR. Even for the images with the highest noise values (*SN R ≈ -* 7.8*dB*) the error in the fitting is less than 0.1*μm*^2^, which is a very small error and constitute a clear evidence that out algorithm have a good performance in the adjustment of the organelles. As expected, the distribution functions of the area error decreases while the level of noise is less in the images, the error in the area approximation is lower (note that when the *SN R* = 3.88*dB* the maximum error obtained in our simulation was around 0.08*μm*^2^). Fig. 4(b) reveals a very small values for the error of the displacement between the center of the organelles. In this case the difference of the distribution functions between the lowest and highest values of SNR is more pronounced than in Fig. 4(a), but for all SNR values the displacement is very small (less than 0.4*μm*).

**Fig 4.**
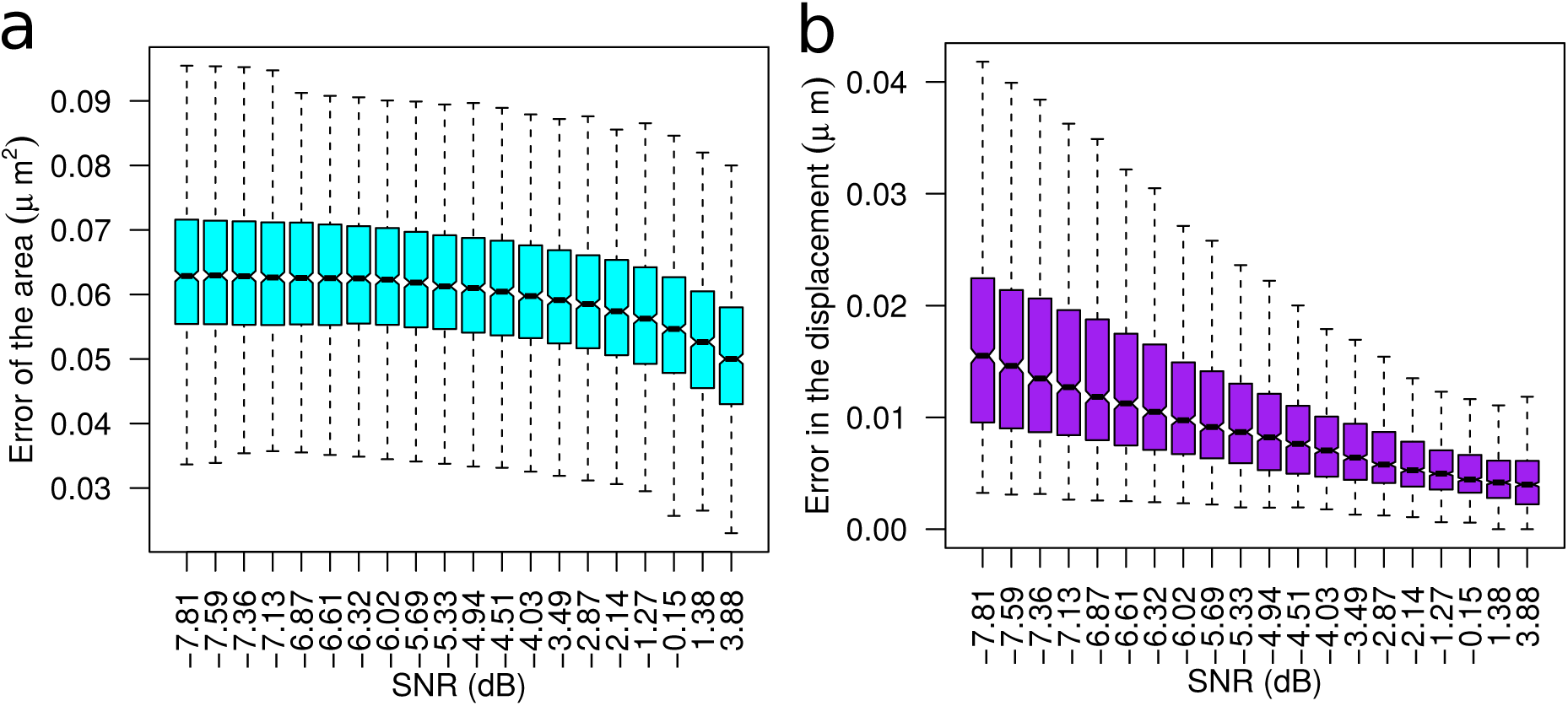
Boxplot graphs for the error in the area and displacement to the center of the “ground truth” organelles. **(a)** Boxplot of the absolute error in the approximation of the organelles’s area (see Note 1). **(b)** Boxplot for the displacement error in the detection and fitting of the organelles. The displacement error was used to establish the criteria of detection and is defined as the Euclidean distance between the center of mass of the organelles *ℒ* and *ℒS*.

Fig. 5 shows the mean values for the area error and for the displacement error as a function of the SNR. In both cases the mean error of the area, and the displacement, decrease when the noise of the image becomes lower. In Fig. 5(a) the upper limit of the confidence interval is always under 0.075*μm*^2^, and in the case of the Fig. 5(b), the displacement error is less than *≈* 0.03*μm* for all the SNR values. These results reflect a very good performance of the algorithm even in images with high noise levels.

**Fig 5.**
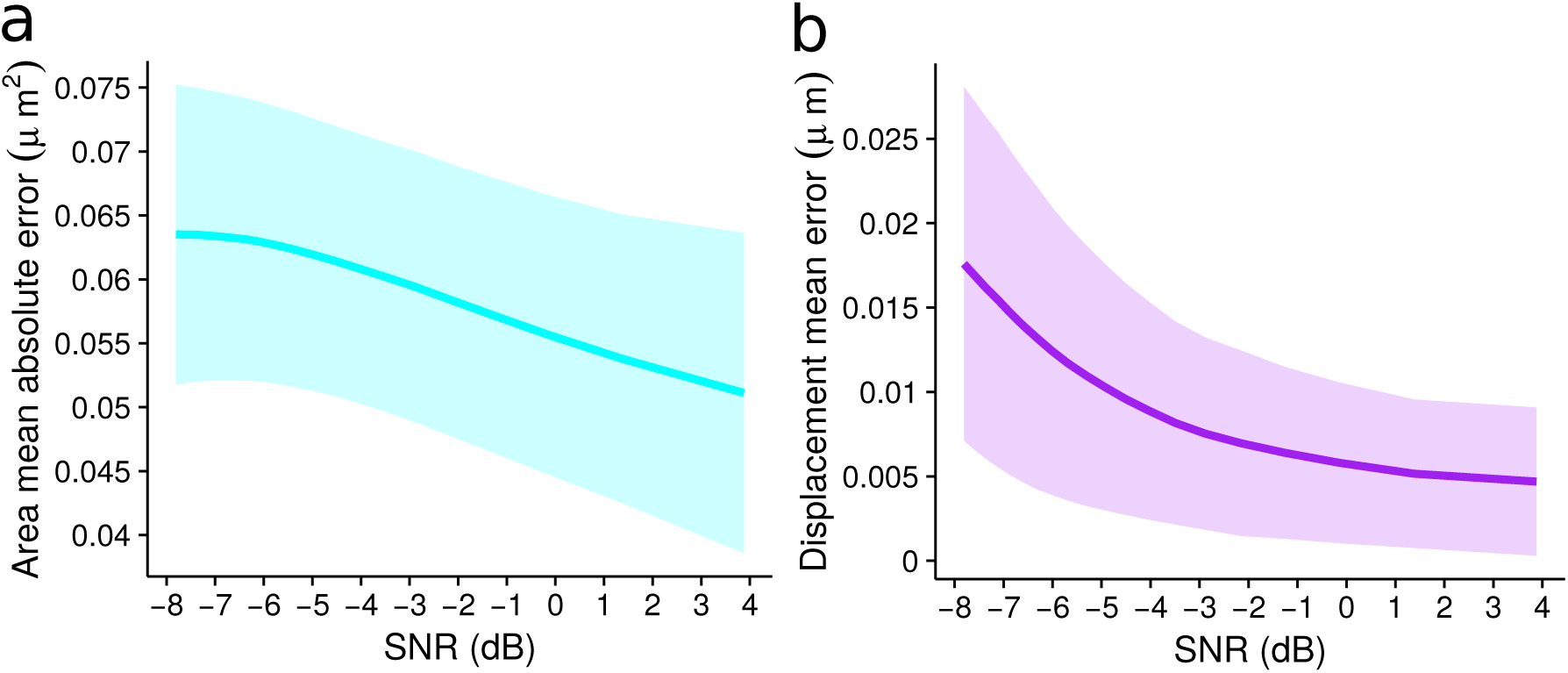
Mean error and confidence interval for the area and displacement of the organelles. **(a)** Mean absolute error for the area between the fitted and simulated organelle. **(b)** Mean of the euclidean distance between the centers of the adjusted and simulated organelle. The confidence interval was computed as a *mean ± sd*, where *sd* is the standard deviation.

### 2.2 Analysis of the algorithm performance on segmenting experimental images

Given the excellent performance of the algorithm on the simulated images, we decided to test it under extreme experimental conditions, i.e., for the detection of cytosolic puncta in presence of a high non-specific fluorescence in the cytosol, achieved by expression of the tandem indicator without special targeting sequence [18, 19].

For our experiments we chose the available mCherry-cpVenus construction, with fluorescent properties and pH-sensitivity close to those of the GFP-RFP tandem indicator used in autophagy studies. HeLa cells transiently expressing the mCherry-cpVenus construction were easily identified by the red-and-yellow-green emission homogeneously distributed throughout the cell cytoplasm, excluding the perinuclear membrane, nucleoli and fine structures most probably corresponding to mitochondria (Fig. 6A).

**Fig 6.**
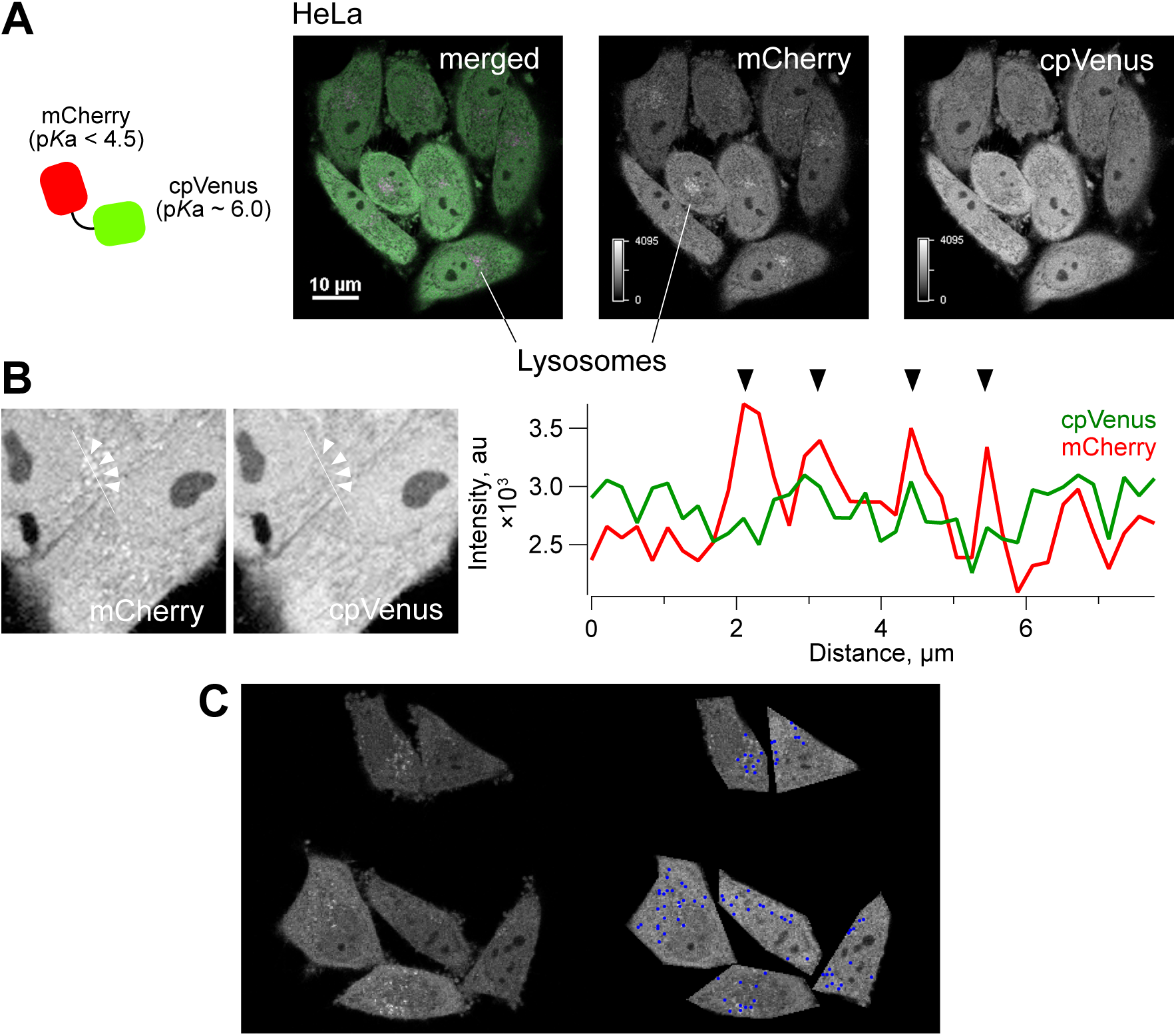
Preliminary testing of the algorithm on experimental images. Given the good performance of the algorithm in segmenting simulated images, we chose to test it for detection of cytosolic puncta on a high cytosolic background in experimental images. For this, we expressed in HeLa a tandem mCherry-cpVenus indicator without targeting sequences and waited 2-3 days until cytosolic punctate structures appeared. A. (left-to-right) Schematic representation of the tandem fluorescent protein expressed in HeLa cells. To the right, merged CLSM image of cells expressing this construction for about 48 hrs, and the fluorescence images of respective mCherry and cpVenus channels are shown. In the mCherry channel and the merged image punctate structures are easily identified, not distinguishable in the cpVenus channel in most cells (B). C. Example of an experimental image of the mCherry channel (left) with cells manually isolated and displaying the centroids positions of the punctate structures identified by the algorithm (blue dots).

Most labeled cells (*∼* 60%, *n* = 89 of 148 analyzed cells) incubated for 48 hrs or longer upon transfection, displayed, in addition to the cytoplasmic fluorescence, dynamic brightly-emitting red puncta, reported previously to correspond to lysosomes [20] (Fig. 6A). As shown in studies by other researchers, the low pKa (*pKa <* 4.5, [21]) and proteolysis-resistance of mCherry, inherited from the parental RFP, for which such properties have been demonstrated [1, 20], was responsible for the appearance of the bright red-emitting cytoplasmic puncta [22]. At the same time, lysosomal sequestration of the mCherry-cpVenus construction resulted in almost complete absence of yellow-green fluorescence in this acidic compartment, as expected from the conditions present in the lumen of these organelles (pH 4.5, [23]) and the relatively high pKa of Venus (pKa = 6.0, [24]), making it sensitive to low pH (Fig. 6B).

We used images obtained in these conditions to test the algorithm performance. An example of manually isolated cells which were further automatically segmented by our algorithm is presented in Fig. 6C. As it can be appreciated, fluorescent puncta are readily identified, even in cells with variable expression of the tandem fluorescent protein, although few spots stayed undetected.

## Conclusion

In the present work we introduced a segmentation algorithm for the analysis of punctate structures similar to those observable in fluorescence images from studies on autophagy. The evaluation of the algorithm segmentation accuracy on more than 20 000 images, simulated using biologically-relevant parameters of puncta number and size, revealed its high robustness even against high background noise levels. This property of the algorithm was confirmed on experimental images, although it came at the expense of some underestimation of the number of punctate structures. We thus suggest that the algorithm could be used to monitor the averaged information from cytosolic puncta, which could be useful in assessing the autophagic flux at a whole cell level, independently of the number of autophagosomes present in the cell.

## Acknowledgments

This work was supported by the Grant-in-aid for Scientific Research on Innovative Areas No. 23115003 “Spying minority in biological phenomena (No. 3306)” from the Ministry of Education, Culture, Sports, Science and Technology of Japan (MEXT) to T.N. YGS received scholarship from CONACyT (Mexico) grant No. 574382.

